# Organic carbon oxidation state shapes fermentative methanogenic microbiomes and controls greenhouse gas fluxes

**DOI:** 10.1101/2025.05.12.653603

**Authors:** Ruiwen Hu, Heidi S. Aronson, Matt E. Weaver, Morgan N. Price, Douglas E. LaRowe, Yuting Liang, Adam M. Deutschbauer, John D. Coates, Hans K. Carlson

**Author notes:** Corresponding author: Dr. Hans K. Carlson Research Scientist, Environmental Genomics and Systems Biology Division E.O. Lawrence Berkeley National Laboratory.

## Abstract

Organic compounds with a negative nominal oxidation state of carbon (NOSC) are thermodynamically recalcitrant in anaerobic ecosystems, but few studies have measured the influence of NOSC on carbon degradation rates, gaseous product yields, or microbiome composition. We amended anaerobic rice paddy sediment microcosms with monomeric organic carbon compounds varying in NOSC. Consistent with thermodynamic and stoichiometric predictions, negative NOSC compounds are catabolized more slowly but produce more methane per mole of carbon. Negative NOSC microbiomes have higher alpha diversity, more syntrophs and methanogens, and fewer fermentative bacteria. Strikingly, fermentative bacterial taxa display genomically encoded NOSC catabolic preferences both in the lab and field. Negative NOSC- preferring fermenters have longer predicted doubling times, consistent with the thermodynamic recalcitrance of their preferred substrates. We propose that microbial NOSC preference can be leveraged for predicting and engineering greenhouse gas fluxes and understanding bacterial population dynamics and trait evolution across redox gradients.

## Introduction

Microbial heterotrophic organic carbon degradation plays a critical role in driving the carbon cycle by transforming photosynthetically fixed carbon into carbon dioxide (CO_2_) and methane (CH_4_). It remains a grand challenge to develop a mechanistic understanding of energy and electron flow from organic carbon through microbiomes to enable optimal management of microbial ecosystems for human and planetary health. There is growing evidence that microbial carbon catabolic pathways are unevenly distributed across microbial genomes, and that organic carbon quantity and quality can be manipulated to drive changes in microbiome composition and element cycling activity in the lab^1–5^ and in the field^6–8^. At the same time, we are improving our ability to link carbon catabolic traits to metabolic niches of microorganisms through genome annotation and laboratory enrichment^9–12^. These advancements, along with improved methods for measuring key organic carbon properties at field scales^13,14^, set the stage for a deeper understanding of microbial carbon transformations.

To generate energy from organic carbon degradation, heterotrophic microorganisms either oxidize organic carbon coupled to reduction of a terminal electron acceptor (e.g. oxygen, nitrate, iron and sulfate respiration) or disproportionate organic carbon (e.g. fermentative methanogenesis to produce CO_2_ and CH_4_) (Fig. 1A). Compared with aerobic respiration, fermentative methanogenesis is much less energetically favorable, and its lower energy yield (Δ*G*) per mole carbon (C) generally leads to lower yields and slower rates of product flux and greater accumulation of intermediates. The electron density of a particular organic molecule also influences the Δ*G* per mole C that can be extracted by microbial catabolism. The electron density per mole C of an organic molecule can be expressed in terms of the average or nominal oxidation state of organic carbon (NOSC) in that molecule^15^. For respiratory metabolism, NOSC predicts the moles of electron acceptor reduced per mole of carbon. For instance, this value corresponds to the chemical oxygen demand (COD) in aerobic respiration. For fermentative methanogenesis, NOSC predicts the CH_4_ and CO_2_ yields^16^, as shown in Fig. 1B and Supplementary Table 6. While these observations follow directly from chemical stoichiometry, to our knowledge, there has been no discussion in the literature on the direct relationship between NOSC and the distribution of methanogenic products.

**Fig. 1.**
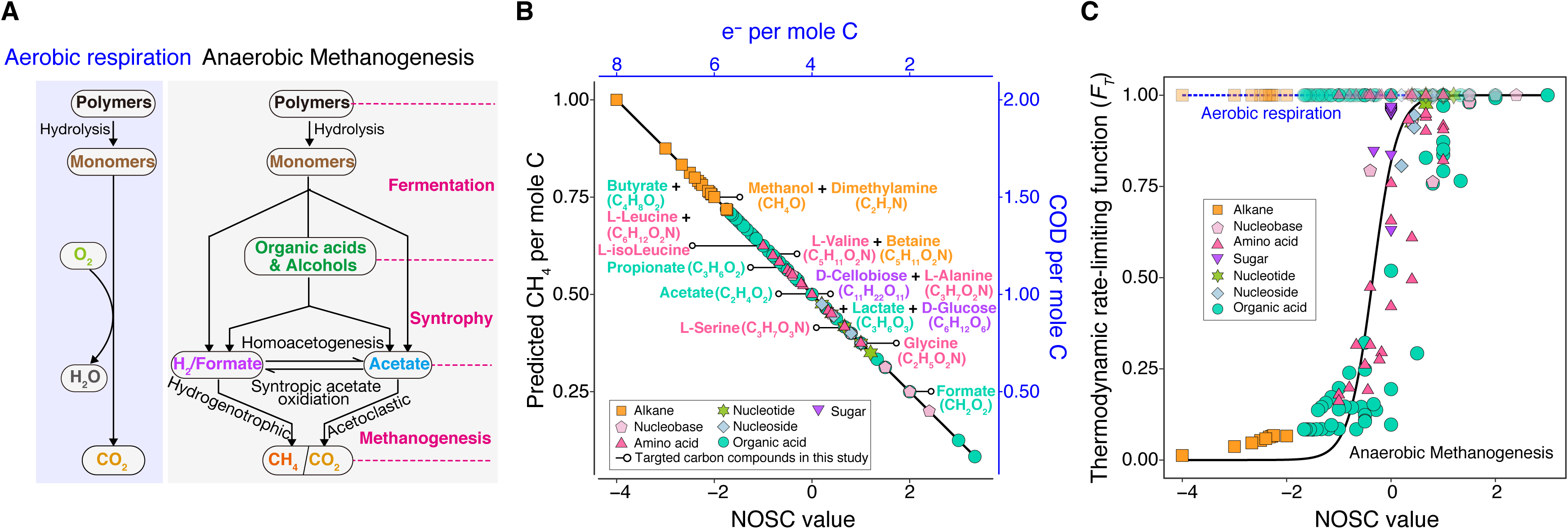
Relationships between nominal oxidation state of organic compounds and thermodynamic recalcitrance and product distribution of microbial metabolism A. Electron flow for heterotrophic carbon degradation coupled to aerobic respiration and fermentative methanogenesis. B. Theoretical methane (CH_4_) production per mole carbon and chemical oxygen demand (COD) per mole carbon (mol O2/mol C) plotted against the nominal oxidation state of carbon (NOSC) and electrons per mole carbon for different organic carbon compounds. C. Relationship between NOSC and thermodynamic rate-liming function (*FT*) for aerobic respiration and anaerobic fermentative methanogenesis.

The greater electron density per mole C in negative NOSC compounds leads to a more favorable Δ *G* per mole C from carbon oxidation coupled to oxygen or nitrate respiration^17^. Thus, aerobic and nitrate respiratory microorganisms may obtain more Δ*G* per mole C, thereby generating more biomass per mole C from oxidation of more reduced organic compounds. However, more reduced organic carbon also requires more energy to be oxidized, which matters more for lower energy metabolisms like sulfate-reduction and methanogenesis (Supplementary Fig. 1)^18^. This thermodynamic recalcitrance is predicted to lead to slower rates of degradation under fermentative conditions where sulfate or carbon dioxide is the sole terminal electron acceptor^15,18^. A thermodynamic rate-limiting term (*F_T_*, see Materials and Methods) has been incorporated into kinetic models to describe how the rates of biologically catalyzed reactions are constrained by their distance from thermodynamic equilibrium—reactions closer to equilibrium proceed more slowly. Specifically, reactions with *ΔG* closer to zero have lower values of the constraint factor *F_T_*compared to more exergonic reactions^18^. When we plot the predicted *F_T_* of Δ *G* per mole electron of the overall metabolic reactions for aerobic respiration and anaerobic fermentative methanogenesis (Fig. 1C), we see a greater thermodynamic recalcitrance (lower *F_T_*) of negative NOSC compounds under methanogenic conditions. In contrast, *F_T_* remains high for all NOSC under aerobic conditions. There is not a perfect correlation between NOSC and *F_T_*, which depends on the Δ *G* per mole electron yield of the metabolism (Fig. 1C, Supplementary Fig. 1), but NOSC is a reasonable proxy that, compared with Δ *G*, is simpler to calculate and measure for complex natural organic carbon using methods like Fourier transform ion cyclotron resonance mass spectrometry (FT-ICR MS), elemental analysis or nuclear magnetic resonance (NMR) spectroscopy^19–21^.

Despite thermodynamic predictions, direct empirical measurements of the degradation rates of organic carbon as a function of NOSC in methanogenic systems remain scarce. Notably, consistent with theoretical expectations, negative NOSC compounds accumulate in anoxic, sulfidic sediments compared to oxic sediments^22–26^. However, negative NOSC compounds are also more hydrophobic and thus less likely to be water-soluble and available for microorganisms^17^. As such, the field studies that have looked at NOSC changes in sediments cannot easily uncouple solubility from microbial energetic controls on organic carbon half-life.

While thermodynamic stability of reduced organic carbon may be beneficial for trapping carbon in soils and sediments, we must also consider its potential climatic consequences. Under methanogenic conditions, negative NOSC compounds yield more CH_4_ per mole C (Fig. 1B), a greenhouse gas with a global warming potential approximately 28 times greater than that of carbon dioxide^27^. Thus, the CH_4_ production rate/yield tradeoffs associated with variations in NOSC, while generally not well-incorporated into field-scale models^6,8,28–32^, are critical for an accurate prediction of greenhouse gas fluxes from methanogenic ecosystems.

The variable thermodynamic recalcitrance of carbon as a function of NOSC across redox clines is also important for understanding the metabolic niche of microorganisms. Oxygen has limited solubility in water and as such water-saturated microbial ecosystems quickly develop hypoxic and anoxic zones. Most prokaryotes are facultative anaerobes and it is estimated that at least 30% of bacteria are capable of fermentative growth^33^. Thus, fermentative bacteria are likely to be very sensitive to organic carbon varying in NOSC and we can postulate that the thermodynamic recalcitrance of reduced organic carbon influences the distribution of carbon catabolic traits in genomes depending on microbial niche and ecological strategy.

In this study we measured the influence of NOSC on the yields and rates of CH_4_ and CO_2_ and microbiome composition in anaerobic fermentative methanogenic laboratory microcosms (Supplementary Fig. 2). We found that more reduced (negative NOSC) carbon yields more CH_4_, but at slower rates. In both laboratory microcosms and field sediments, reduced carbon is associated with greater alpha diversity and greater relative abundance of syntrophs and methanogens. We also found distinct genomically encoded preferences for NOSC among fermentative bacteria that predict their abundances in both laboratory microcosms and natural field sediments. Fermentative bacteria that prefer thermodynamically recalcitrant, reduced carbon substrates have longer predicted minimal doubling times based on the codon bias in their ribosomal protein genes. Our results shed light on the ecological dynamics of fermentative bacteria and provide insights into the co-evolution of carbon catabolism with other microbial traits. More broadly, we propose that microbial NOSC preference serves as a predictive framework for estimating greenhouse gas flux potential based on microbiome composition, guiding targeted interventions to regulate carbon fluxes and fermentation end-products in methanogenic ecosystems, and elucidating the evolutionary dynamics of microbial metabolic traits.

## Results

### Carbon oxidation state controls greenhouse gas production yields and rates in rice field sediments

To measure the influence of carbon oxidation state on fermentative methanogenic microbiomes we amended rice field sediment microcosms with organic carbon compounds spanning NOSC from −2 to +2 (Materials and Methods). We selected compounds based on (i) prevalence in rice field systems (cellobiose and glucose), (ii) importance as fermentation intermediates (propionate, butyrate and lactate), (iii) ability to serve as direct substrates for methanogenesis (acetate, formate, methanol and dimethylamine) and finally, (iv) a consistent structural class that spans a wide range of NOSC (amino acids). Apart from single carbon compound microcosms, we also tested amino acid mixtures combined in equal proportions (L-leucine/L-serine, L-isoleucine/L-serine, glycine/L-serine and L-leucine/L-isoleucine). After 190 days, microcosms amended with more reduced carbon (NOSC < 0) produced significantly more CH_4_ than CO_2_, while more oxidized carbon (NOSC > 0) yielded more CO_2_ than CH_4_ (Fig. 2A, Supplementary Table 1 and Fig. 3). For example, L-leucine (NOSC = −1) yielded 1.75 times more CH_4_ (0.564 ± 0.01 mM/mol C) than L-serine (NOSC = +0.67) (Fig. 2A), which is consistent with theoretical predictions. We observed a significant negative correlation between CH_4_ yields and NOSC in for all compounds (R = −0.76, *P* < 2.2e−16), as well as within the amino acids (R = −0.93, *P* < 2.2e−16) and or organic acids (R = −0.62, *P* =0.00019) (Fig. 2B).

**Figure 2.**
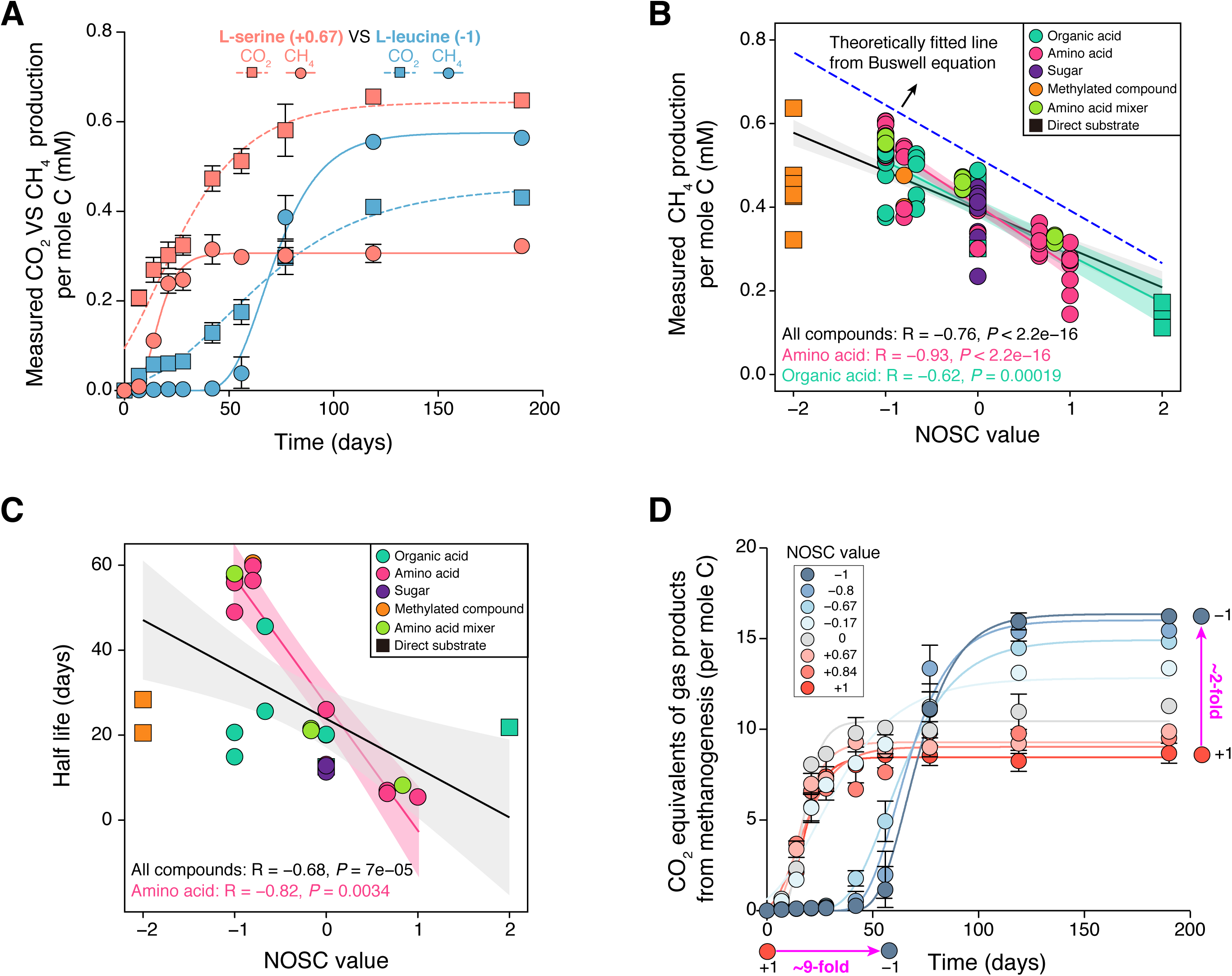
Influence of monomeric organic carbon varying in NOSC on methane and carbon dioxide production rates and yields in methanogenic microcosms from rice field sediment. A. Methane (CH_4_) and carbon dioxide (CO_2_) concentrations per mole of substrate carbon over 190 days in rice field microcosms amended with L- serine (NOSC = +0.67) and L-leucine (NOSC = −1). B. NOSC of different monomeric organic carbon compounds from different chemical classes plotted against measured CH_4_ yields per mole C after 190-days in rice sediment microcosms. Colors indicate compound class: Organic acids, teal (butyrate, propionate, lactate, acetate and formate); amino acids, pink (L-leucine, L-isoleucine, L-valine, L-alanine, glycine and L-serine); sugars, purple (cellobiose and glucose); methylated compounds, orange (dimethylamine, methanol and betaine); mixtures of amino acids, light green (L- leucine/L-serine, L-isoleucine/L-serine, L-serine/glycine, L-leucine/L-isoleucine); Circles indicate fermentable carbon compounds and boxes represent direct methanogenic substrates (formate, acetate, methanol and dimethylamine). The dashed line indicates theoretical methane yields for fermentative methanogenesis and solid lines represent Spearman’s rank correlations with shaded 95% confidence intervals. Spearman’s rho and p-values (*P* < 0.05 was considered statistically significant) are shown for the set of all compounds (black), or the sets of amino acids (red) or organic acids (teal). C. NOSC of different monomeric carbon compounds plotted against measured half-life of CH_4_ production in days. Coloring and shapes are the same as panel B. Solid lines represent Spearman’s rank correlations with shaded 95% confidence intervals. Spearman’s rho and p-values are shown for the set of all compounds (black), or the sets of amino acids (red). D. CO_2_ equivalent concentration per mole substrate carbon (per mole substrate CH_4_ * 28 + per mole substrate CO_2_) in rice field sediment microcosms amended with individual amino acids or mixtures varying in NOSC from −1 to +1 over 190-day. Arrows indicate the fold increase in CO_2_ equivalents and half-life when NOSC is varied from −1 to +1.

**Fig. 3.**
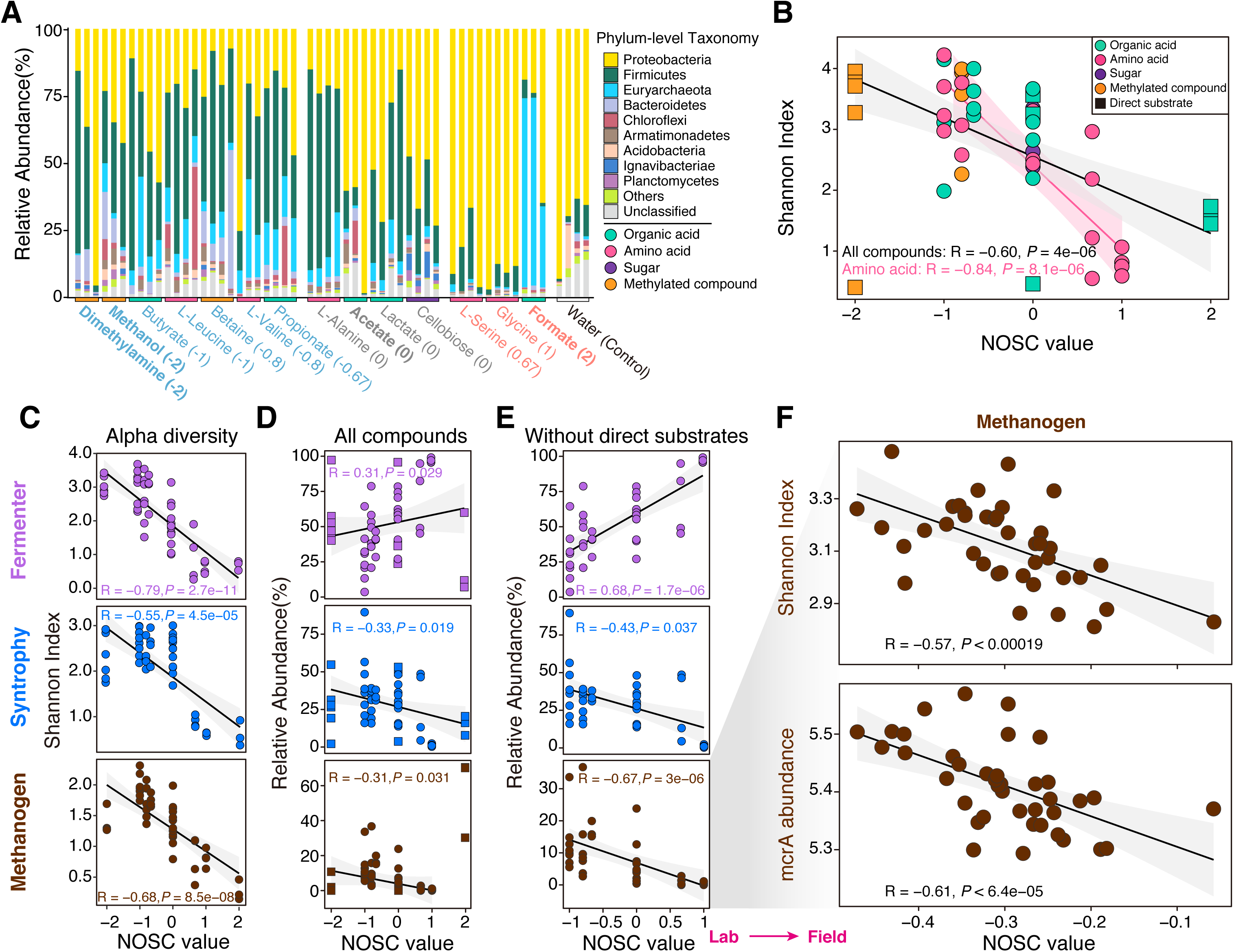
Changes in methanogenic microbiomes in response to variations in NOSC. A. Phylum-level relative abundances from 16S rDNA amplicon sequencing after 190 days in methanogenic microcosms amended with different organic carbon compounds with NOSC from −2 to +2. B. NOSC of organic carbon compounds plotted against the Shannon index from 16S rDNA amplicon sequencing in rice field sediment microcosms after 190 days. Symbols and colors are the same as Figure 2B and 2C. C. NOSC of organic carbon compounds plotted against the Shannon index in microcosms for sub- populations of fermentative bacteria (purple), obligate and facultative syntrophs (blue) and methanogenic archaea (brown) (See Materials and Methods for details on selecting sub-populations). D. NOSC of organic carbon compounds plotted against the relative abundances in microcosms of fermentative bacteria (purple), syntrophs (blue) and methanogenic archaea (brown) for the set of all organic carbon compounds. E. NOSC of organic carbon compounds plotted against the relative abundances in microcosms of fermentative bacteria, syntrophs and methanogenic archaea for the subset of carbon substrates excluding the direct substrates of methanogens (formate, acetate, methanol and dimethylamine). F. Shannon index of the methanogenic archaeal microbiome based on archaeal specific 16S rDNA amplicon sequencing and the relative abundance of the *mcrA* gene from amplicon sequencing of rice paddy field sediments plotted against the NOSC of soil organic carbon. Spearman’s rho and p-values (*P* < 0.05 was considered statistically significant) are shown for all correlations.

Our observed CH_4_ yields are consistent with theoretical predictions for both individual carbon compounds and compound mixtures^16^ (Fig. 1). However, we noticed that across all conditions our microcosms produced ∼0.378 mM on average less CH_4_ than predicted (Fig. 2B). This slight discrepancy is likely due to consumption of organic carbon electrons coupled to other respiratory metabolisms such as the trace ferric iron present in the rice field sediment. Regardless, the majority of organic carbon electron equivalents can be accounted for based on organic carbon flow through fermentation, syntrophy and methanogenesis to produce CO_2_ and CH_4_ at the predicted ratios^16^ (Fig. 1).

In addition to controlling greenhouse gas yields, NOSC is expected to control the thermodynamic favorability and thus influence the rate of organic carbon mineralization under anaerobic conditions^15^ and some field studies have observed enrichment of reduced organic carbon in anoxic sediments^25^. However, to the best of our knowledge, the half-life of organic carbon varying in NOSC under methanogenic conditions has not been measured in the laboratory. We observed slower rates of gas production and growth with more reduced organic carbon compounds compared with more oxidized organic carbon compounds (Fig. 2C). For example, L-serine was rapidly degraded by methanogenic microbiomes to produce CH_4_ and CO_2_ while the half-life of L-leucine was much longer (Fig. 2A).

Across all carbon compounds we tested, we observed a negative correlation between half-life and NOSC. This correlation was better when we omitted direct methanogenic substrates and when we focused only on amino acids (R = −0.82, *P* = 0.0034) (Fig. 2C) Also, we observed a less robust correlation between NOSC and half-life for the set of organic acids that were tested. In reality, thermodynamic inhibition (*F_T_*) is dependent on the Δ *G* per mole electron of the overall methanogenic metabolic reaction for each compound. Thus, some compounds are relatively more thermodynamically recalcitrant than would be predicted based on their NOSC (Fig. 1).

For example, while propionate has a more positive NOSC than butyrate, its complete oxidation reaction to CO_2_ is more endergonic than that of butyrate (Fig. 1C). Consistent with the differences in their *F_T_*, we observed a longer half-life for propionate than for butyrate.

CH_4_ is a 28 times stronger greenhouse gas than CO ^27^, so we can also convert the gas concentrations we measure to CO_2_ equivalents. When comparing amino acids that vary in NOSC from −1 to +1 we observe a ∼2-fold increase in climate warming potential across this range of NOSC (Fig. 2D) with a ∼9-fold longer half-life. Thus, while more reduced organic carbon is slower to be degraded it has a greater climate warming potential when it is degraded under methanogenic conditions.

### Carbon oxidation state shapes the diversity and composition of methanogenic microbiomes

We measured the taxonomic composition of the methanogenic microcosms using 16S rDNA amplicon sequencing (Fig. 3A). At the phylum level, positive NOSC microcosms are dominated by *Proteobacteria* (*Pseudomonadota*) (average abundance 87.62%). The one exception is formate which is a direct substrate for methanogens and enriches for *Euryarchaeota*. In contrast, negative NOSC microcosms are dominated by *Firmicutes* (39.98%) and *Euryarchaeota* (10.76%). For organic carbon with NOSC = 0, microcosms have intermediate enrichment of these dominant taxa (Fig. 3A). We also observed higher alpha diversity (Shannon index) in more negative NOSC microcosms (Fig. 3B). Across all microcosms, more reduced carbon is correlated with more diverse microbiomes. This is true both for the set of all compounds (R = −0.60, *P* < 4e−06) and the subset of amino acids (R = −0.84, *P* < 8.1e−06).

We next classified all ASVs (amplicon sequence variants) as fermentative bacteria (fermenters), syntrophs or methanogens based on their taxonomy (Supplementary Table 2) to analyze how carbon oxidation state influenced each of these sub-populations in the methanogenic microcosms. As with the set of all ASVs, we observed increased diversity within sub-group in more negative NOSC compounds (Fig. 3C). However, the relative abundance of each sub-group changed with NOSC; more methanogens and syntrophs and fewer fermenters are found in negative NOSC compounds (Fig. 3D). Notably, when we exclude direct methanogenic substrates (e.g. formate, acetate) and focus only on the set of fermentable carbon substrates we observed significant correlations between NOSC and the relative abundances of methanogens, syntrophs and fermenters (R_Fermenter_ = 0.68, *P* = 1.7e−06; R_syntroph_ = −0.43, *P* = 0.037; R_methanogen_= −0.67, *P* = 3e−06) (Fig. 3E).

To support the generality of our observed relationships between NOSC, alpha diversity and relative abundance of methanogens, we turned to field datasets. In a previous study on rice paddy fields, Shannon diversity of methanogens and the abundance of the *mcrA* gene were quantified^13^ and carbon composition measurements by NMR^34^ enabled us to calculate NOSC (Supplementary Fig. 4) in field samples. In rice field sediments, we found significant correlations both between Shannon diversity and NOSC, and between *mcrA* abundance and NOSC (Fig. 3E and 3F). Notably, we did not observe a difference in the relative proportion of hydrogenotrophic versus acetoclastic methanogens. This is consistent with the reported complex relationships between these two functional groups and the role of other factors like temperature in influencing the relative contributions of acetate versus formate versus hydrogen as methanogenic substrates in natural soils and sediments^35^.

### Carbon oxidation state selectively enriches for distinct populations of fermentative bacteria

To more deeply characterize how NOSC influences gene content in fermentative methanogenic microbiomes we selected replicate rice sediment microcosms amended with five different amino acids that vary in NOSC from −1 to +1 (L-leucine, L-valine, L-alanine, L-serine and glycine) and sequenced gDNA to generate metagenome assembled genomes (MAGs) (Fig. 4). A total of 312 high-quality or medium-quality MAGs were recovered from these five amino acid enrichments, consisting of 267 MAGs affiliated with 13 bacterial phyla and 45 MAGs affiliated with 3 archaeal phyla (Supplementary Fig. 5A and Table 3). Based on their taxonomy and annotations of carbon catabolic traits (Supplementary Table 4), we divided these 312 MAGs into fermentative (145 MAGs), syntrophic (65 MAGs), methanogenic (45 MAGs) and unknown (57 MAGs) groups. As in our amplicon data, we observed significant correlations between NOSC and the relative abundances of fermenters, syntrophs and methanogens (R_Fermenter_ = 0.74, *P* = 0.00022; R_syntroph_ = −0.97, *P* = 2.4e−12; R_methanogen_= −0.91, *P* = 3.4e−08) (Supplementary Fig. 5B).

**Fig. 4.**
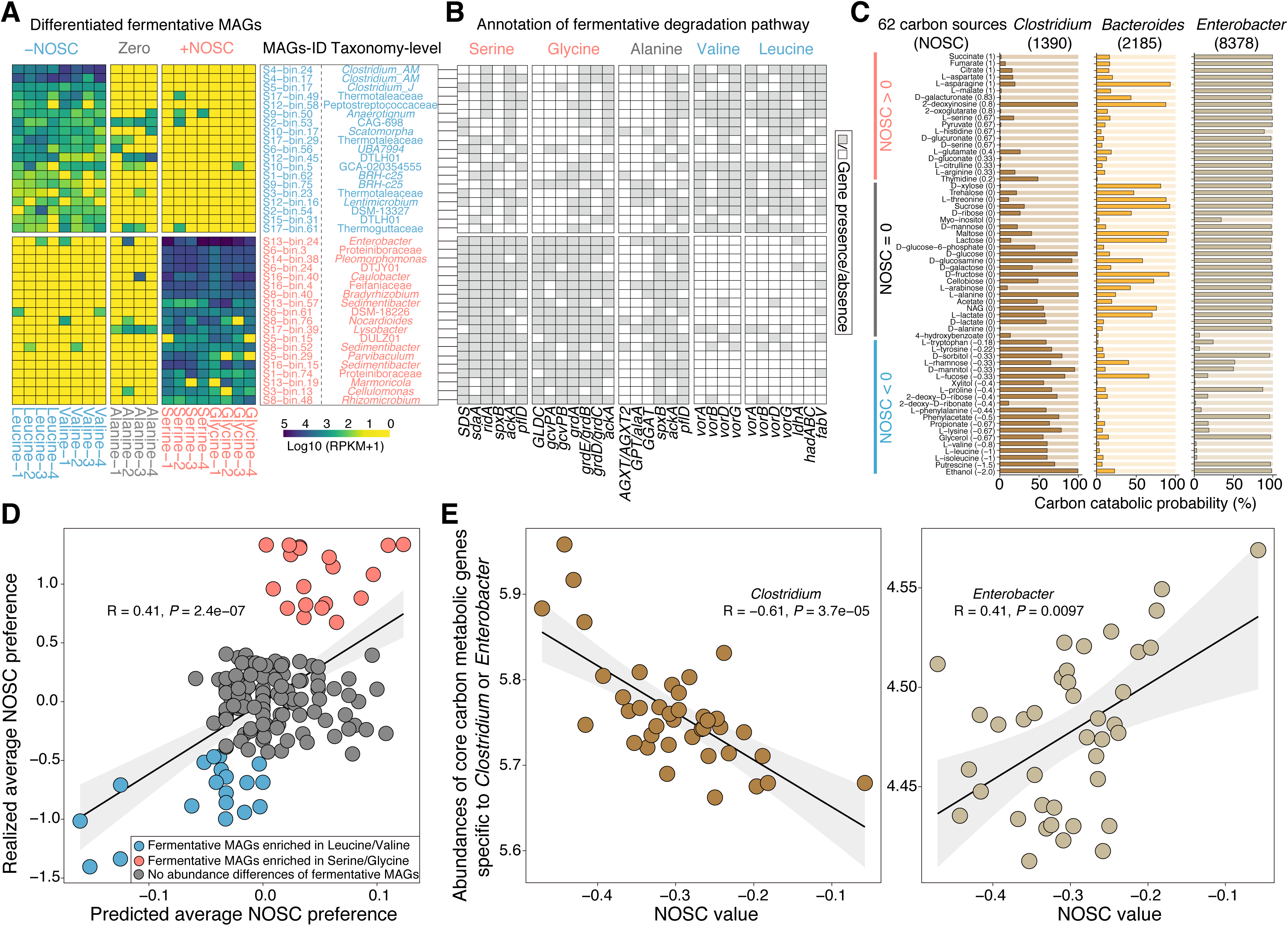
Influence of five amino acids varying in NOSC on fermentative microbiome composition and gene content. A. Relative abundances of fermentative bacterial metagenome-assembled genomes (MAGs) differentially enriched in replicate microcosms amended with either negative NOSC amino acids (L-leucine and L-valine) or positive NOSC amino acids (L-serine and glycine). Each group of positive and negative NOSC enriched MAGs is sorted based on average Log2 fold-change (Log2FC). Mags-ID and Taxonomy-ID are given for each bin (See Supplemental Table S3). B. Presence or absence in differentially enriched MAGs (Panel A) of genes involved in fermentative metabolism of five amino acids. C. Gapmind predictions of catabolic probabilities for 62 organic carbon compounds sorted by NOSC from −2 to +1 in genomes from the genera *Bacteroides* (2,185), *Clostridium* (1,390) and *Enterobacter* (8,378) from AllTheBacteria database^77^. D. Predicted average NOSC preference from GapMind plotted against measured average NOSC preference of each fermentative MAG across all amino acid enrichments. Solid line indicates Spearman rank correlation. Points are colored based on their relative differential enrichment in positive or negative NOSC amino acids (Panel A). E. Abundance of core carbon metabolic genes from fermentative bacteria (genera *Clostridium* and *Enterobacter*) in rice paddy field soils based on GeoChip 5.0 microarray plotted against soil organic matter NOSC (see reference^13,77^ and materials and methods).

We used differential abundance analyses (Materials and Methods) to identify MAGs significantly differentially enriched in L-serine and glycine (positive NOSC) microcosms relative to L-leucine and L-valine microcosms (negative NOSC) and found fermentative bacterial MAGs exhibited the most pronounced differentiation (Fig. 4A and Supplementary Fig. 5C). The dominant fermenter MAGs enriched in L-serine and glycine include representatives of the genera *Enterobacter*, *Lysobacter*, *Nocardioides* and *Sedimentibacter*, while fermenter MAGs enriched in L-leucine and L-valine enrichments are *Clostridium_AM, Clostridum_J*, *UBA7994*, *Lentimicrobium* and *BRH- c25*.

To understand the basis for their differential selection, we searched each fermenter MAG for genes known to be involved in catabolism of the five amino acids added to the microcosms (Fig. 4B). Strikingly, fermenters enriched in L-serine and glycine microcosms have serine ammonia-lyase (*SDS*), serine dehydratase (*sdaA*), glycine reductase (*GLDC/gcvPAB*) and glycine dehydrogenase (*grdABCDE*) but lack many genes necessary for utilization of L-leucine and L-valine. In contrast, fermenters enriched in L-leucine and L-valine microcosms have 2-ketoisovalerate ferredoxin reductase (*vorABCD*) and 4-methyl-2-oxopentanoate reductase (*idhA*), but lack genes required for catabolism of L-serine and glycine (Fig. 4B). To further support this genomic prediction we isolated representative dominant fermenters from L-serine and L-leucine microcosms. We found that the *Enterobacter* sp. isolate from L-serine microcosms could ferment L-serine and glycine, but not L-leucine or L-valine. In contrast, the *Clostridium* sp. isolate could ferment L-leucine and L-valine but not L-serine or glycine (Supplementary Fig. 6). Taken together, these results strongly suggest that the fermentative bacterial populations in the microcosms were selectively enriched based on their carbon catabolic preferences for amino acids varying in NOSC.

We next sought to determine if the carbon catabolic preferences of the dominant fermentative bacteria (*Enterobacter*, *Clostridium* and *Bacteroides*) in our rice field microcosms are representative of the prevalence of these traits within each genus. We used a recently developed computational tool, GapMind^10^ to annotate catabolic pathways for 62 monomeric organic compounds in genomes from across the genera *Enterobacter, Clostridium,* and *Bacteroides* (Fig. 4C). GapMind annotations indicate that *Enterobacter* tend to possess catabolic pathways for positive NOSC compounds. In contrast, *Clostridium* tend to possess catabolic pathways for negative NOSC compounds and *Bacteroides* tend to have pathways for compounds with NOSC close to zero (mostly sugars) (Fig. 4C).

For each fermenter MAG in our microcosms, we calculated a predicted average NOSC preference based on GapMind annotations of its genome. We also calculated the realized average NOSC preference of each fermentative MAG by weighting the NOSC of carbon compounds in a given microcosm by the MAG’s relative abundance. A significant correlation was observed between predicted and realized NOSC preferences across all fermentative MAGs in our microcosms (R =0.41, *P* =2.4e-7) (Fig. 4D). Based on this observation, we hypothesized that NOSC preference could influence composition of the fermentative microbiome across gradients of NOSC in the field. We again turned to the published rice paddy field dataset where we could compare NOSC with other microbiome traits^13,34^. The authors of this previous study used a DNA hybridization based approach and specific gene probes for core metabolic enzymes specific to either *Enterobacter* or *Clostridium.* The abundance of these genus-specific enzymes serves as a reasonable proxy for the relative abundance of these taxonomic groups in field samples. Consistent with our hypothesis, we found more *Enterobacter* specific genes measured at more positive NOSC and more *Clostridium* specific genes at more negative NOSC (Fig. 4E).

### Fermentative bacteria have carbon oxidation state preferences

Based on the observation that the fermentative bacteria in our microcosms and the *Enterobacter* and *Clostridium* in rice paddies display NOSC preferences, we were encouraged to systematically investigate NOSC preference across a broader range of fermentative bacteria. A recent study identified a set of bacterial genera that have been phenotypically confirmed to possess fermentative growth capabilities^33^. We retrieved genomes representative of each genus known to be capable of fermentation from GTDB and used GapMind to predict carbon catabolic preferences for these fermentative genera, calculated the probability that a representative of a given genus will utilize a given organic carbon compound and calculate their average NOSC preferences.

The complete dataset of all genera with any characterized fermentative representative is in the Supplementary data (Supplementary Table 5) but we focused on the set of genera where more than half of the isolates are experimentally shown to be capable of fermentation and rank ordered these genera based on their average NOSC preference (Fig. 5A). Genera with the greatest positive NOSC preference include *Enterobacter, Pantoea, Erwinia, Staphlococcus, Serratia, Haemophilus,* and *Vibrio.* Genera with the greatest negative NOSC preference include *Clostridium, Azospirillum, Rhodoferax, Ruminoclostridium, Fibrobacter, Ruminococcus, Spiroplasma,* and *Listeria.* Of the genera with realized positive-NOSC preferences in our microcosms (Fig. 4A), our results are very consistent with predictions from GapMind. For example, we found positive NOSC-preferences for *Enterobacter, Rhizomicrobium*, *Sedimentibacter*, *Caulobacter*, and *Lysobacter* and strong negative NOSC preferences for *Clostridia, Peptostreptococcus* and *Thermogutta* (Supplementary Table 5). Taken together, these results suggest that NOSC preferences can be predicted based on microbial gene content and that this trait is mostly conserved at the genus level.

**Fig. 5.**
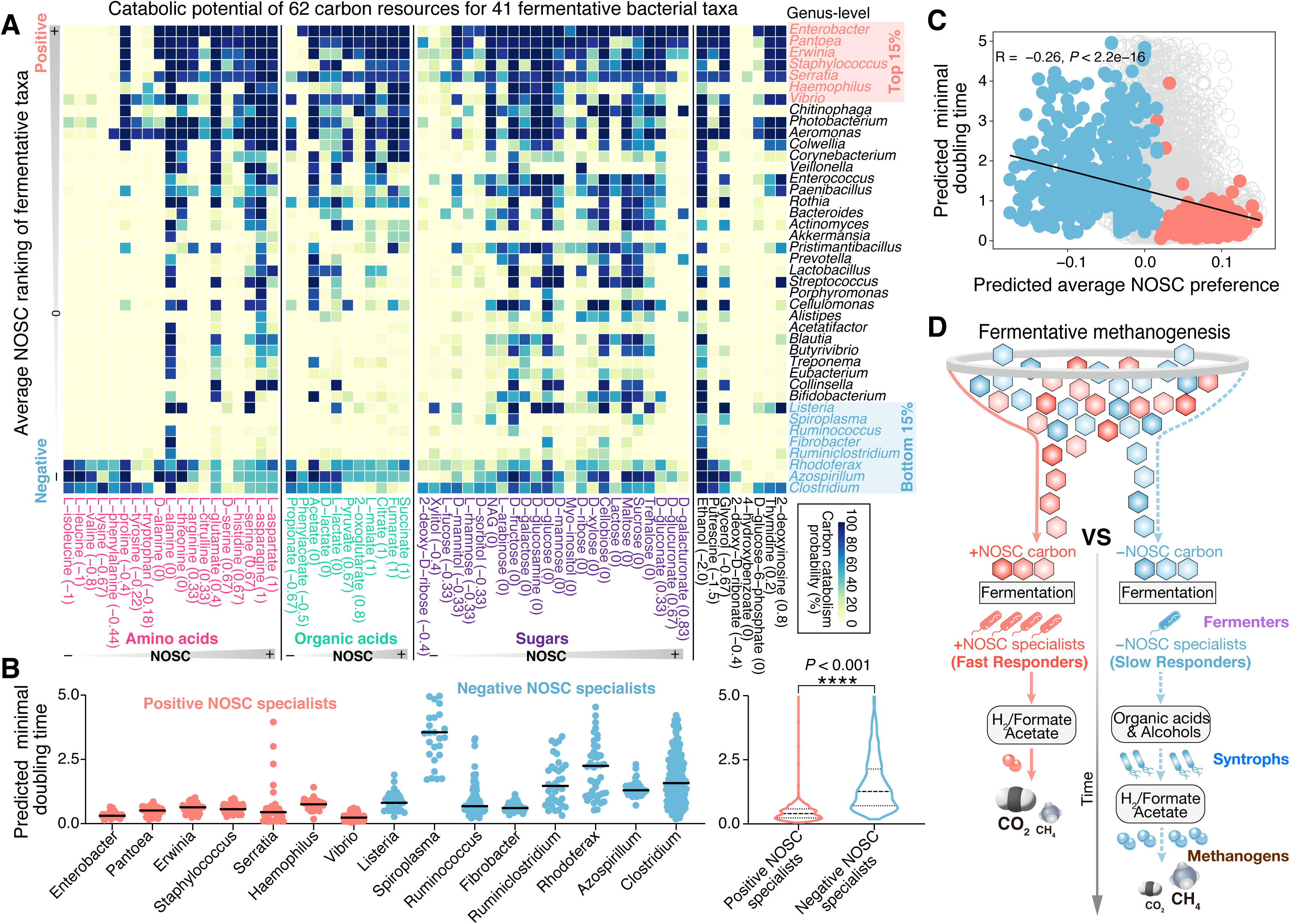
NOSC preferences of fermentative bacteria. A. GapMind predictions of carbon source utilization from Genome Taxonomy Database (GTDB) genomes representing known fermentative genera^33^. Heatmap shows the probability that a given carbon source is catabolized by a representative of each genus (with more than 30 genomes). Genera are sorted based on their average NOSC preference across 62 organic carbon compounds with NOSC from −2 to +1. A total of 7,159 genomes were collected from the GTDB. B. Comparison of predicted growth doubling time between negative NOSC and positive NOSC fermentative specialists. C. Spearman correlations between the predicted growth doubling time of 7,159 fermentative genomes and their predicted average NOSC preference across 62 organic carbon compounds. Negative and positive NOSC specialists were colored by sky blue and pink. D. Concept model describing the impact of NOSC on the fermentative methanogenesis.

Given that carbon compounds with negative NOSC degrade more slowly under fermentative methanogenic conditions (Fig. 2), we hypothesized that fermenters specializing in negative NOSC substrates may also exhibit slower growth rates, reflected in longer minimal doubling times. To test this hypothesis, we calculated the predicted minimal doubling time based on ribosomal codon bias ^36,37^ for bacterial genomes from genera with positive and negative NOSC preferences. Consistent with our hypothesis, positive NOSC specialists (Top 15%) tend to have shorter predicted doubling times compared to negative NOSC specialists (Bottom 15%) (Fig. 5B).

Furthermore, we observed a significant correlation (R = -0.26, *P* <2.2e-16) between average NOSC preference and doubling time across all genomes from genera in which more than half of the known isolates have demonstrated fermentative capabilities (Fig. 5C). The correlation between predicted NOSC preference and minimal doubling time is weaker for genera with lower experimentally confirmed fermentation capacity or for non-fermentative genera (Supplementary Fig. 7), however the Spearman’s correlation is still significant suggesting that NOSC preference and growth rate are likely coupled traits for all heterotrophic bacteria. Overall, our results support the hypothesis that the thermodynamics of anaerobic growth shape carbon catabolic preferences in bacteria.

## Discussion

We used a laboratory cultivation-based approach to measure the influence of NOSC on fermentative methanogenesis, including its impact on the yields and production rates of greenhouse gases (CH_4_ and CO_2_) and the associated microbiome composition and gene content. Consistent with thermodynamic and stoichiometric predictions, organic compounds with negative NOSC were degraded slower, but yield more CH_4_ per mole C. Negative NOSC microbiomes exhibit greater alpha diversity, an increased abundance of syntrophs and methanogens, and distinct populations of fermentative bacteria both in laboratory microcosms and in rice paddy fields (Fig. 5D). To our knowledge, these relationships between carbon oxidation state and methanogenic microbial ecology have not been observed but have important implications for our understanding of controls on greenhouse gas fluxes and anaerobic microbiome composition. Additionally, our demonstration that NOSC preference is a genomically predictable and taxonomically conserved microbial trait will help in mechanistic interpretation of microbiome composition and enable better prediction and control of microbiomes.

While many factors may influence the distribution of organic carbon catabolic traits across microbial genomes, we found that many genera of fermentative bacteria have conserved preferences for negative NOSC or positive NOSC compounds based on gene annotations. This prediction is derived from gene content analysis, yet numerous pathways and transporters involved in organic carbon metabolism remain unidentified and challenging to annotate. However, our findings, in conjunction with previous literature^38–40^, support the hypothesis that NOSC preference represents an underappreciated microbial trait with significant implications for shaping microbiome composition in environmental ecosystems.

We found that *Enterobacter* generally prefer positive NOSC amino acids (e.g., L-serine and glycine), while *Clostridium* tend to prefer negative NOSC amino acids (e.g., L-leucine, L-valine and L-isoleucine) both in our microcosms and rice field samples. Although few studies have explicitly investigated the role of carbon selectivity in shaping microbiome composition, a previous study demonstrated that L-serine is strongly selective for *Enterobacter* in the human gut^41^. Additionally, other field studies suggest that soil microbiota composition varies with NOSC^38^, and that different metagenomes and bacterial proteomes vary in their oxidation state^39,40^. Our results suggest that NOSC preferences likely explain these previous observations. Bacteria with a preference for negative NOSC compounds tend to exhibit weaker codon bias for ribosomal protein genes, which correlates with longer predicted doubling times (Fig. 5C). Although multiple factors could influence both NOSC preference and doubling time, our results suggest that NOSC preference and doubling time likely co-evolve. In dynamic anaerobic microbial ecosystems receiving inputs of complex detrital carbon with varying NOSC, faster-growing fermenters are likely to utilize more thermodynamically labile positive NOSC compounds, while slower-growing fermenters are more likely to encounter higher concentrations of negative NOSC compounds. Recent advances in measuring *in situ* microbial growth rates have highlighted the importance of this trait as critical in determining microbiome dynamics in response to variable nutrient levels and water saturation^37,42^.

Our study provides insights into the functional ecology of methanogenic ecosystems. Microbial methanogenesis is responsible for a significant portion of the increase in methane emissions^43–45^. Consequently, for developing effective mitigation strategies for the climate-warming effects driven by methanogenesis, it is critical to systematically identify the key factors controlling this process. Temperature, pH, and salinity are well-established controls on the mechanisms and fluxes of methane^35,46–48^. However, due to the complexity of overlapping environmental factors and the role of methane oxidation in regulating methane fluxes, it has been challenging to directly observe the impact of organic carbon composition on CH_4_/CO_2_ ratios in field settings^49,50^. Nonetheless, the elemental stoichiometries of organic carbon compounds are clearly linked to these ratios^16^, and we have connected this relationship to the concept of carbon oxidation state. This framework allows us to demonstrate the rate-yield trade-offs inherent in varying carbon oxidation states in methanogenic systems. Despite the importance of carbon oxidation state, it is rarely discussed in studies focused on the thermodynamic preservation of organic carbon in wetland soils and sediments, although it has critical implications for land management strategies^30,51,52^.

Here we focused on NOSC preference as an important microbial trait because, compared with Δ *G*, NOSC is simpler to calculate measured in natural systems. However, thermodynamic recalcitrance of organic carbon under methanogenic conditions is more likely to be controlled by Δ *G* of methanogenesis and the related concept of the thermodynamic rate limiting function (*FT*). Ultimately, we anticipate that calculation of a Δ*G* preference and an associated power (P = Δ*G* x Δt) preference^53^ will be useful for understanding the environmental niche of heterotrophic bacteria. Interrogating how these microbial traits co-occur with other traits related to anabolism and stress tolerance will help us develop a microbial physiology based view of the tree of life. Future research also should explore the role of NOSC in conjunction with other factors that influence methane fluxes, such as methylotrophic methanogenesis, polyphenol degradation, or alternative electron acceptors. Additionally, there is a potential to use NOSC preferences in microorganisms to predict how NOSC will reliably alter microbiomes by enriching fermentative populations that differ in their sensitivity to inhibitors. We are optimistic that a deeper mechanistic understanding of methane flux controls will ultimately enable more effective management of both natural and engineered ecosystems.

## Materials and methods

### Thermodynamics calculations for organic carbon compounds varying in nominal oxidation state

For a set of 141 diverse organic compounds we calculated the nominal oxidation state of carbon (NOSC), CH_4_:CO_2_ ratios, chemical oxygen demand per mol C, the Gibbs energy of carbon oxidation, *ΔG_cox_*, the Gibbs free energy of microbial respiratory metabolisms with different terminal electron acceptors, and the thermodynamic rate limiting function (*F_T_*) for these reactions (Fig. 1 and Supplementary Table 6). NOSC was calculated using Equation 1, where *Z* is the net charge of the compound, and *a, b, c, d, e*, and *f* correspond to the stoichiometric numbers of the elements C, H, N, O, P, and S^15^. CH_4_:CO_2_ ratios were calculated using the Buswell Equation, where the amount of methane produced by the fermentation of an organic compound is predicted by Eq. 2 and the amount of CO_2_ produced by this fermentation reaction is predicted by Eq. 3. For both Eq. 2 and Eq. 3, *a, b, c, d*, and *f* correspond to the stoichiometric numbers of the elements C, H, N, O, and S.

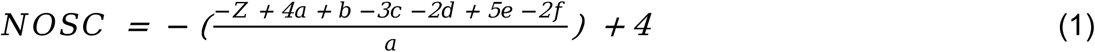

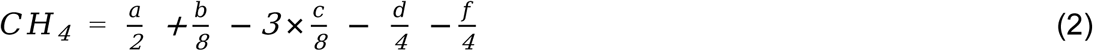

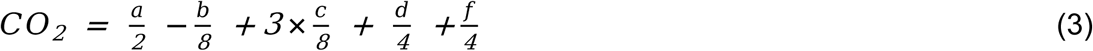

The Gibbs energy of reaction, *ΔG_r_*, was calculated for carbon oxidation half reactions coupled to several terminal electron accepting half reactions, including methanogenesis, sulfate reduction, denitrification and aerobic respiration, using

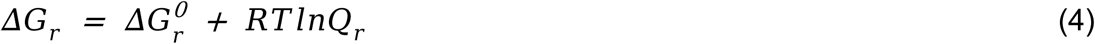

where Δ*G^0^_r_* is the standard state Gibbs energy, *R* represents the universal gas constant, and *T* refers to the temperature in Kelvin, and *Q*_r_ denotes the reaction quotient. Values of Δ*G^0^_r_* were calculated for each reaction at 25°C and 1 bar with the revised Helgeson-Kirkham-Flowers (HKF) equations of state^54–56^ using the “subcrt” command from the R software package CHNOSZ v1.4.1^57^. Thermodynamic data in CHNOSZ are derived from the OrganoBioGeoTherm database, which come from a number of sources, as documented in (https://chnosz.net/download/refs.html). Values of Q_r_ were calculated with

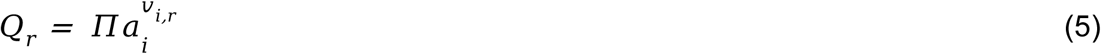

where *a_i_* corresponds to the activity of the *i* th species raised to its stoichiometric coefficient, *v_i,r_*, in the *r*^th^ reaction, which is positive for products and negative for reactants. For all reactions, the pH was set at 7, and the activities of all carbon substrates were set at 10^-3^, which is approximately equivalent to 1 mM. Environmentally-relevant activities of electron acceptors and products were chosen, with bicarbonate activity set at 2x10^-3^ (≈ 2 mM), sulfate activity set at 2.8x10^-2^ (≈ 28 mM), oxygen activity set at 5.18x10^-4^ (≈0.518 mM), methane activity set at 10^-4^ (≈100 µM), N2 concentration set at 6x10^-4^ (≈600 µM), and nitrate, bisulfide, and ammonium activities set at 10^-4^ (≈100 µM).

*F_T_* was calculated for each reaction at 25°C with the function

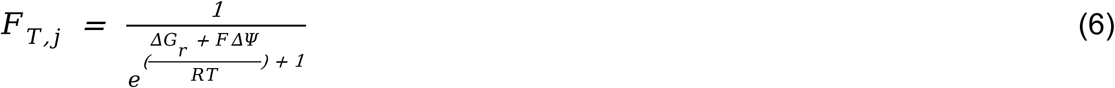

where *ΔG_r_* is the Gibbs energy of the reaction, calculated with Eq. 4, *F* is the Faraday constant (96.485 KJ/mol), *Δψ* is the membrane potential (set at 80, 120, and 240 mV), *R* represents the universal gas constant, and *T* refers to the temperature in Kelvin^18^.

Microcosms from rice field sediment cultivation and monitoring Microcosms were prepared from rice paddy sediment collected from the California Rice Experiment Field Station at 955 Butte City Hwy, Biggs, CA (39.46 °N, 121.73 °W). Each microcosm in a 22.5 mL anaerobic Balch tube received ∼5 grams of sediment and 10 mL of an anaerobic chemically defined basal medium (Supplementary Table 7). Individual organic carbon compounds were added from concentrated stock solutions at concentrations indicated in Supplementary Table 1 and Table 2. All chemicals are from Sigma-Aldrich (St Louis, Mo, USA). Anaerobic batch tube microcosms were maintained at 30°C in an incubator. We also amended 1 mL raw cultures in 96 deepwell blocks (Costar) with dilute sediment suspensions (∼0.05 g sediment per culture) in an anaerobic chamber (Coy). Inoculated microplates were sealed with silicon microplate seals (VWR), sealed in anaerobic boxes (BD) with a BD GasPack scrubber and paper towel soaked in concentrated 100 mM sodium sulfide and incubated at 30°C in an incubator in an anaerobic chamber (Coy).

Headspace CH_4_ and CO_2_ gas concentrations in anaerobic Balch tube microcosms were measured with a Agilent gas chromatograph model 7890A (G344OA) with a Superlco SP™-2380 Fused Silica Capillary column containing a proprietary, bonded silica-based stationary phase. CH_4_ was measured using a flame ionization detector (FID) and CO_2_ was measured with a thermal conductivity detector (TCD). The GC oven temperature was held at 50 °C for 60 minutes to allow for non-carrier gases to elute. The FID heater was kept at 300°C; air flow was 400mL/min; hydrogen gas fuel flow was 30mL/min; and nitrogen gas flow was 25 mL/min. (H_2_ and N_2_ are the carrier gases for FID). The TCD was kept at 300 °C; reference flow was at 30 mL/min; and the makeup flow (He- the carrier gas) was kept at 2 mL/min. Methane and carbon dioxide standard gases (Scotty Analyzed Gases) were used for calibration. GC ChemStation (Agilent Technologies) computer software was used for quantification. For microplate enrichments growth was measured by sub-sampling 100 μL of culture into a shallow 96 well plate (Greiner) in an anaerobic chamber using an Aviden Micropro 200 pipettor (Rainin) and measuring optical density (OD_600_) with a Tecan M1000 Pro microplate reader (Tecan Group Ltd Männendorf, Switzerland).

### DNA extraction and 16s rDNA amplicon analysis

For 16S rDNA amplicons, gDNA extractions from microcosms were performed with slight modifications from a previous method^58^. 1 mL of each microcosm was removed with a syringe and needle and transferred into a 96 deep well block (Costar) and centrifuged to collect biomass (4000 rcf, 10 min). Pelleted cells were resuspended in 100 µL sterile water and 180 µL enzymatic lysis buffer (ELB) which contains 20 mM Tris-HCl pH 8.0, 1 mM sodium EDTA (TE) with 1.2% Triton X-100 was added. 20 µL lyzosyme (25 mg/mL) was added and the samples incubated overnight at 37°C. The following morning 20 µL of proteinase K (20 mg/mL) and 200 µL 4 M guanidinium HCL were added and the samples incubated for 2 hrs at 55°C. Next, 4 µL RNAse A (100 mg/mL) was added and samples incubated for 2 h at room temperature. 300 µL of 100% ethanol was added to precipitate DNA and samples were then applied to 96 well silica membrane plates (BPI-Tech, San Diego, CA, USA). DNA bound to membranes was washed twice with 400 µL 70% ethanol:TE and eluted in 100 µL TE. An Avidien Micropro 200 (Avidien) was used to transfer large volumes of buffers and multichannel pipettes (Rainin) used to apply samples to 96 well columns and add low volume reagents. A vacuum manifold (Qiagen, Redwood City, CA, USA) was used to perform column purification steps. Extractions using this method were found to be similar in terms of DNA yield and recovery to using the Gene QIAamp 96 DNA QIAcube HT Kit (Qiagen).

gDNA template was added to a PCR reaction to amplify the V4/V5 16S gene region using the 515F/ 926R primers based on the Earth Microbiome Project primers^59,60^, but with in-line dual Illumina indexes^11,61^. The amplicons were sequenced on an Illumina MiSeq (Illumina, San Diego, CA, USA) with 2×300 bp Ilumina v3 reagents. Reads were processed with custom Perl scripts implementing PEAR for read merging^62^, USearch^63^ for filtering reads with more than one expected error, demultiplexing using inline indexes and Unoise^64^ for filtering rare reads and chimeras. 16S sequences in the relative abundance table were searched against the RDP database^65^ to assign taxonomy.

### DNA extraction, metagenomic sequencing and data analysis

For metagenomic sequencing, 5 mL from each microcosm was sampled and centrifuged (4000 rcf, 10 min) to collect biomass. The Power Soil DNA Isolation Kit (Mo Bio Laboratories, Carlsbad, California, USA) was used for gDNA extraction according to manufacturer instructions. DNA concentration and quality were assessed by NanoDrop ND-2000 Spectrophotometer (Thermo Fisher Scientific, MA, USA) and agarose gel electrophoresis. ∼1.00 μg of DNA was prepared for metagenomic libraries using a NEBNext Ultra II DNA Library Prep Kit (New England Biolabs, MA, USA) according to the manufacturer’s instructions. The DNA quality was determined using a LabChip GX Touch HT (PerkinElmer, USA) and quantified by real-time quantitative PCR. Samples were sequenced individually with NovaSeq 6000 (Illumina, San Diego, CA, USA), resulting in 2 × 150 bp paired-end reads with an average of 16.57 Gbp per sample (Supplementary Table 8). Metagenomic raw reads were trimmed using FastP (v0.23.4)^66^ with default parameters to remove low-quality reads, adapters, and polyG sequences. We then used the metaWRAP pipeline for genome assembly and binning^67^. The sequences combined by trimmed reads of each enrichment culture from 5 amino acids were assembled with MEGAHIT (options: −mink 21 −maxk 141 −step 12) to generate contigs^68^. We then performed genome binning of assembled reads by using MetaBAT2 (version 2.12.1)^69^, CONCOCT (version 0.4.1)^70^, and MaxBin2 (version 2.2.4)^71^ based on tetranucleotide frequencies, scaffold coverage, and GC content. Finally, we further improved the resulting bins to obtain the final metagenome-assembled genomes (MAGs) set via metaWRAP’s Bin_refinement module and Reassemble_bins module. The quality of MAGs was assessed with CheckM2 v1.0.2^72^ against GTDB database (release 207v2), while MAGs with <50% completeness or >10% contamination were removed. All MAGs were finally dereplicated using dRep (version 3.4.5)^73^, where an average nucleotide identity (ANI) of 95% was used as cutoff to dereplicate MAGs from the secondary ANI comparison. Meanwhile, coverage value of each MAG was expressed as RPKM (Reads Per Kilobase of transcript per Million mapped reads) and calculated using CoverM v0.6.1 genome mode (https://github.com/wwood/CoverM). The taxonomic assignments of MAGs were conducted using GTDB-Tk v2.3.2^74^. The phylogenomic tree of these MAGs was constructed with GToTree v1.8.4^75^ setting parameters (-*H* Bacteria_and_Archaea -G 0.5). To classify the 312 MAGs into fermenter, syntroph, and methanogen groups, initial functional grouping was performed based on their taxonomic information and assigned uncertain MAGs to the unknown group. Completeness of their carbon metabolism profiles across 62 carbon compounds was predicted from the categories of organic acids, amino acids, and sugars (Supplementary Table 4) using GapMind for carbon metabolism annotations^10^ and we manually refined the initial classification by defining the syntroph group as organisms that preferentially utilize organic acids while having limited capacity to metabolize fermentable sugars. By differential abundance analysis of MAGs from fermenter, syntroph, and methanogen groups, 38 fermentative MAGs, 24 syntrophic MAGs and 30 methanogenic MAGs were identified as significantly enriched in negative NOSC microcosms (L-leucine/L-valine) or positive NOSC microcosms (L-serine/glycine) (Figure 4A and Supplementary Fig. 5C). Fermentation pathways in each fermentative MAG were annotated (Supplementary Table 9) based on the curated Metacyc database^76^ and DRAM v1.5.0^9^. Moreover, the average NOSC preference of each fermentative MAG was quantified by calculating the mean NOSC of all 62 carbon compounds predicted to be utilized by GapMind. The realized average NOSC preference of each fermentative MAG was assessed by calculating the mean abundance of each MAG across all microcosms multiplied by the NOSC value of each corresponding microcosm.

### Physiological assays of fermentative bacterial isolates

For isolation of fermentative bacterial isolates, 1 mL of each microcosm was plated onto anoxic solid blood agar (BD). Single colonies were re-streaked three times for further purification and then single colonies were transferred into anaerobic chemically defined medium with glucose as the sole fermentable carbon source. All fermentative bacterial isolates were cryopreserved in 25% glycerol and stored at −80 °C and gDNA was extracted as above for full length 16S rDNA Sanger sequencing (27F/1492R). To determine each isolate’s capacity to ferment L-leucine, L-valine, L-alanine, L-serine and glycine we transferred washed cells into basal medium containing one of the five amino acids (10 mM) and measured optical density (OD_600_) over time using a Spectronic 20D spectrophotometer (Supplementary Fig. 6).

### Gapmind annotations for carbon metabolism and maximal growth rates prediction of fermentative bacteria in AllTheBacteria and GTDB database

To investigate the NOSC-dependent carbon catabolic preferences from the three fermentative genera (*Bacteroides*, *Clostridium* and *Enterobacter*) that dominated our microcosms, 11,953 genomes were collected from these genera, including 2,185 *Bacteroides* genomes, 1,390 *Clostridium* genomes and 8,378 *Enterobacter* genomes, in AllTheBacteria database with 661K high-quality bacterial genomes^77^. Using GapMind for carbon metabolism annotations^10^, the potential of each genome to utilize 62 carbon compounds was predicted. To determine whether a carbon source can be utilized by bacteria from the *Bacteroides*, *Clostridium*, and *Enterobacter* genera, the GapMind annotation results for each genome must meet two criteria: (i) The total score of the carbon catabolic pathway must be greater than 0, where each high-confidence step scores is +1, each medium-confidence step scores is −0.1, and each low-confidence step scores is −2. (ii) The carbon catabolic path has no low-confidence steps. The number of *Bacteroides*, *Clostridium*, and *Enterobacter* bacteria capable of utilizing each carbon source in GapMind annotations was predicted and their proportions in these three genera as the genus specific carbon catabolic probability were quantified (Fig. 4C).

To further investigate the NOSC-dependent carbon catabolic preference across fermentative bacteria, 24,175 bacterial genomes were collected from the publicly available Genome Taxonomy Database (GTDB) based on their membership in a set of bacterial genera that have been phenotypically confirmed to be capable of fermentative growth in a recent study^33^. By analyzing fermentation probabilities (the proportions of bacteria with phenotypically confirmed fermentative capabilities in each bacterial genus) of the recorded bacterial genera in this study, these GTDB bacterial genomes were classified into three distinct groups: the high-probability fermentative bacterial group (7,159 genomes belonging to 1,030 bacterial genera with fermentation probability ≥ 50%), the low-probability fermentative bacterial group (3,598 genomes belonging to 166 bacterial genera with 0 < fermentation probability < 50%), and the non-fermentative bacterial group (13,418 genomes belonging to 1,488 bacterial genera with fermentation probability = 0) (Supplementary Table 5). Using GapMind, the carbon catabolic potential for 62 carbon compounds in these 24,175 bacterial genomes was predicted and their predicted average NOSC preference was quantified. The number of utilized carbon compounds for each genome and carbon catabolic probability of these genomes at genus-level were calculated (Fig. 5A and Supplementary Table 5). Based on distributions of the predicted average NOSC preference across all genomes from the high-probability fermentative bacterial groups (Supplementary Fig. 8), we identified bacterial genera with average NOSC preference in the top 15% as positive NOSC specialists and bottom 15% as negative NOSC specialists.

We used the R package gRodon (v2.0)^78^ to predict the minimal doubling times at the genome- and genus-level of 24,175 bacterial genomes belonging to bacterial genera from the high-probability, the low-probability and the non-fermentative bacterial groups (Figure 5B). The genes encoding ribosomal proteins, which are typically highly expressed in bacteria, were identified by hmmsearch^79^ against bacterial ribosomal proteins (TIGR01632) in the TIGRFAMs database^80^ with trusted cutoff (parameter “-- cut_tc”). To increase the comparability of the predicted minimal doubling time across various bacteria, we removed data with a predicted minimal doubling time greater than 5, as a predicted minimal doubling time of around 5 already corresponds to no selection for efficient translation of ribosomal proteins^36^. Spearman’s correlation was used to identify the relationship between the predicted minimal doubling time and the predicted average NOSC preference of 24,175 bacterial genomes from the high-probability fermentative, the low-probability fermentative and the non-fermentative bacterial groups (Fig. 5C and Supplementary Fig. 7).

### Distribution patterns of NOSC and methanogens in soil organic matter (SOM) from 39 typical paddy fields

Using previous soil organic matter (SOM), GeoChip 5.0 microarray, and archaeal specific 16S rDNA amplicon sequencing datasets from 39 rice paddy fields across northern to southern China^13,34^, the relationships between NOSC of SOM and diversity or abundances of soil methanogens, as well as abundances of specific fermentative taxa *Enterobacter* or *Clostridium* were investigated. Based on the relative abundances of functional groups, including alkyl C (0-45 ppm), methoxyl C (45–60 ppm), O-alkyl C (60–95 ppm), Di-O-alkyl C (95–110 ppm), aromatic C (110–145 ppm), phenolic C (145– 160 ppm), and carbonyl C (160–220 ppm) and their corresponding reference NOSC values^19,20^, NOSC values of SOM in each rice paddy soil sample were calculated through weighted summation (Supplementary Fig. 4). The gene *mcrA* encoding the methyl coenzyme M reductase enzyme was used to estimate the methanogenic capacity of microbial communities^81^ All *mcrA*-related probes detected in more than half of the replicates from the same sampling site in the GeoChip 5.0 microarray dataset were selected. Abundance of the *mcrA* gene in each soil sample was calculated by normalizing to the total intensity of the detected probes, multiplying by a constant, followed by natural logarithm transformation. Similarly, we quantified the abundances of carbon degradation-related probes from the GeoChip 5.0 microarray dataset specific to genera *Enterobacter* and *Clostridium* in each soil sample. The abundance of genus-specific genes is a reasonable proxy for the relative abundance of these taxa in these soil samples^82^. The archaeal specific 16S rRNA sequencing was conducted using the primer pair 1106F and 1378R^83^. Spearman’s correlation was used to identify the relationship between NOSC of SOM and relative abundances of *mcrA* gene, or relative abundances of genera *Enterobacter* and *Clostridium*. All details regarding soil sampling, the NMR measurements of SOM, the GeoChip 5.0 microarray experiments and 16S rRNA amplicon sequencing were described in these previous studies^13,34^.

## Statistical Analysis

All statistical analyses and plotting were conducted using GraphPad Prism (version 8.0) and R statistical language (version 4.02). CH_4_ and CO_2_ production curves were plotted and fitted using the Gompertz Model in GraphPad Prism software. Differential abundance analyses of fermentative MAGs, syntrophic MAGs and methanogenic MAGs between negative NOSC microcosms (L-leucine/L-valine) and positive NOSC microcosms (L-serine/glycine) were performed using the R package DESeq2. Wilcoxon signed-rank test (a significance level set at *P* < 0.05) was used to explore the differences of maximal growth rates of fermentative bacteria between positive NOSC specialists and negative NOSC specialists. Ridgeline plots of predicted average NOSC preference distribution of 24,175 genomes from the high-probability fermentative group, the low-probability fermentative group and the non-fermentative group were visualized by the R package ggridges. All correlation analyses were evaluated by a linear fit and Spearman test using the R packages ggpubr and ggplot2. Heatmap plots were visualized by pheatmap and viridis packages. The maximal growth rates of fermentative bacteria were estimated by the R package gRodon (v2.0).

## Acknowledgements

This study was financially supported by the Energy & Biosciences Institute (EBI) through the EBI-Shell program and some co-authors are members of ENIGMA (Ecosystems and Networks Integrated with Genes and Molecular Assemblies; (http://enigma.lbl.gov), a Science Focus Area Program at Lawrence Berkeley National Laboratory, US Department of Energy, Office of Science, Biological and Environmental Research Program under contract number DE-AC02-05CH11231 to Lawrence Berkeley National Laboratory.

## Author contribution

W.H. and H.K.C. conceived and designed the study. R.W.H., M.E.W. and H.K.C. performed the laboratory work and detailed the sampling. R.W.H., H.S.A., M.N.P., D.E.L. and H.K.C. carried out the bioinformatics and statistical analysis, and thermodynamic calculations. R.W.H. and H.K.C. wrote the first draft of the manuscript. R.W.H., H.S.A., M.E.W., M.N.P., D.E.L., Y.T.L., A.M.D., J.D.C. and H.K.C. discussed results and edited. All authors read and approved the final version of the manuscript.

## Data availability

All data needed to evaluate the conclusions in the paper are present in the paper and/or the Supplementary Information. The nucleotide sequences for 16S rRNA gene amplicon sequencing and metagenomic sequencing were deposited in the SRA database under accession numbers PRJNA1188626 and PRJNA1187553, respectively.

## Code availability

All software packages utilized in this study are publicly accessible, and no original code is reported in this study.

