## Supplementary Information for "Organic carbon oxidation state shapes fermentative methanogenic microbiomes and controls greenhouse gas fluxes"

*Corresponding author:

Number of figures.: 8

**Supplementary Tables**

**Table S1**: Time-series measurements of greenhouse gases of our enrichments cultured on 15 carbon sources, including 11 single carbon compounds and 4 mixed amino acids.

**Table S2**: 16S rDNA relative abundance and taxonomy, organic carbon compound metadata and end-point methane measurements for rice paddy field microcosms.

**Table S3**: 312 Metagenome-assembled genomes (MAGs) and their abundances acquired in five amino acid enrichments with NOSC from −1 to +1.

**Table S4**: Gapmind annotations of 312 Metagenome-assembled genomes (MAGs) acquired in five amino acid enrichments with NOSC from −1 to +1.

**Table S5**: Carbon catabolic probabilities of 24,715 genomes from GTDB database at genus level for Gapmind 62 organic carbon compounds.

**Table S6**: Thermodynamics parameters for 141 organic carbon sources varying in nominal oxidation state from −4 to +3.

**Table S7**: Chemically defined media for microcosms from rice field sediments.

**Table S8**: The output of metagenomic raw data of five amino acids with four replicates.

**Table S9**: The fermentation pathways of five amino acids in the curated Metacyc database.

**Supplementary figures**

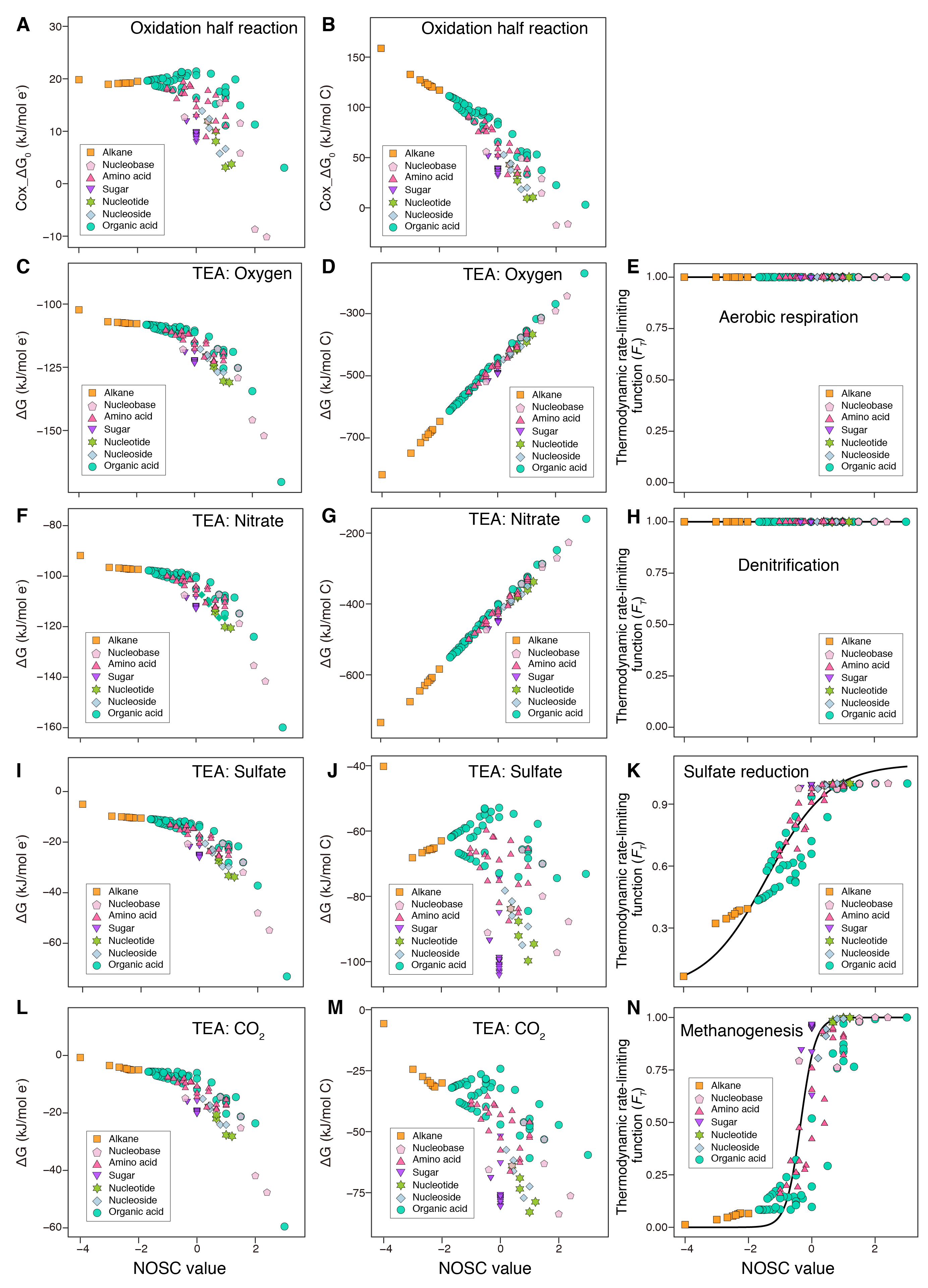

**Figure S1**: Relationships between nominal oxidation state of organic compounds (NOSC) and the energy$\Delta$G per mole electron (e^-^) and per mole carbon (C) for the oxidative half reaction and different terminal electron acceptors (TEA). **A.** the energy$\Delta$G per mole electron (e^-^) for Carbon oxidation half reaction. **B.** the energy$\Delta$G per mole carbon (C) for Carbon oxidation half reaction. **C.** the energy$\Delta$G per mole electron (e^-^) for Aerobic respiration. **D.** the energy$\Delta$G per mole carbon (C) for Aerobic respiration. **F.** the energy$\Delta$G per mole electron (e^-^) for Denitrification. **G.** the energy$\Delta$G per mole carbon (C) for Denitrification. **I.** the energy$\Delta$G per mole electron (e^-^) for Sulfate reduction. **J.** the energy$\Delta$G per mole carbon (C) for Sulfate reduction. **L.** the energy$\Delta$G per mole electron (e^-^) for Methanogenesis. **M.** the energy$\Delta$G per mole carbon (C) for Methanogenesis. For various TEAs: **E., H., K. and** **N.,** the relationship between NOSC and the thermodynamic rate-liming function (*F_T_*) is also shown.

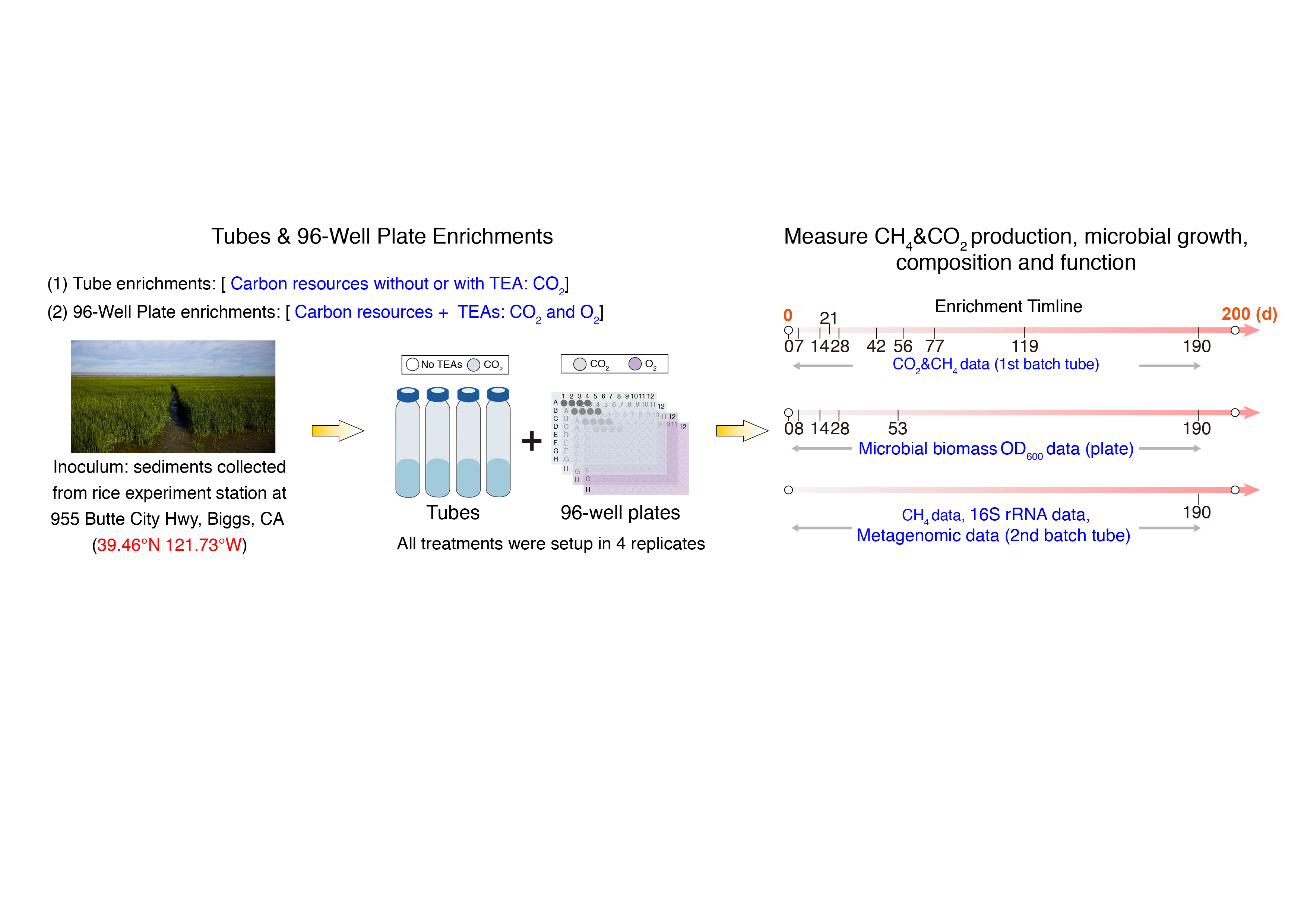

**Figure S2**: Workflow for studying relationships between nominal oxidation state of organic compounds and fermentative methanogenesis via anaerobic tube and plate enrichment cultures and measurement of gas products, microbial growth, microbial community composition and function over 190 days. Terminal Electron Acceptors (TEAs).

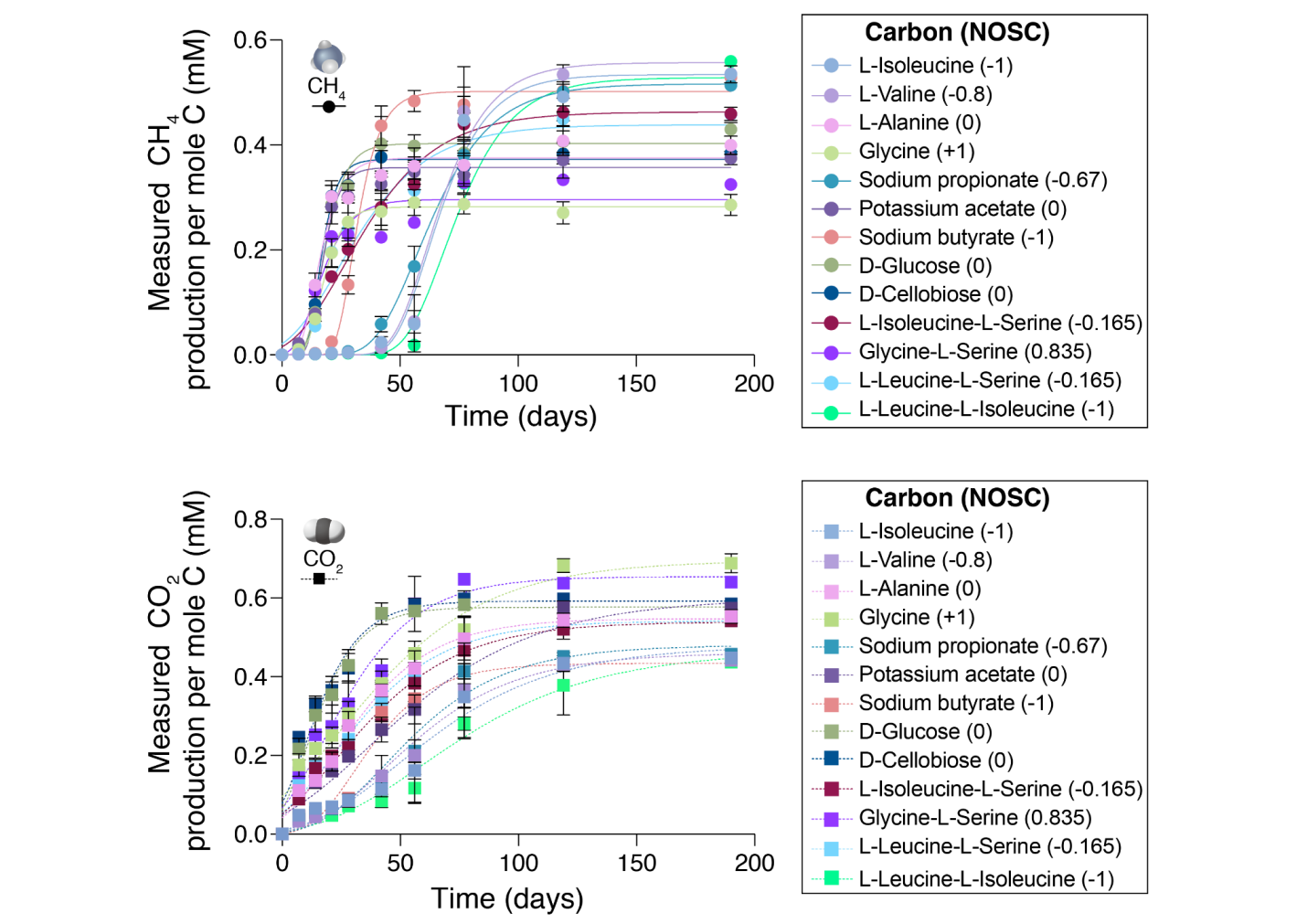

**Figure S3**: Measurements of methane (CH_4_) and carbon dioxide (CO_2_) concentrations per mole of substrate carbon over 190 days in rice field microcosms amended with different monomeric organic carbon sources varying in NOSC.

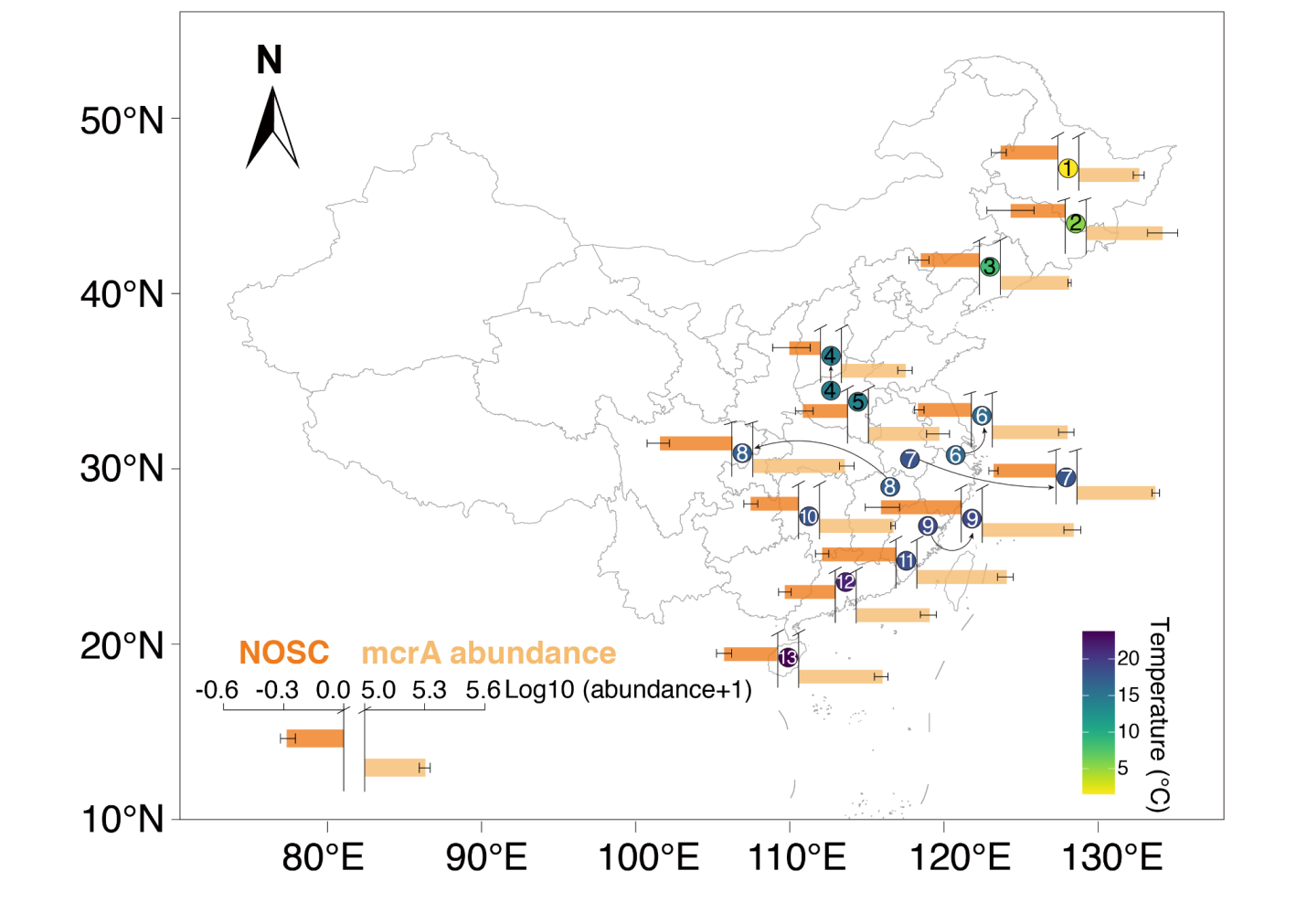

**Figure S4**: Distributions of nominal oxidation state of carbon (NOSC) in soil organic matter (SOM) and functional gene abundances of *mcrA* (the core gene in the methanogenic pathway) from 39 typical paddy fields across northern to southern China.

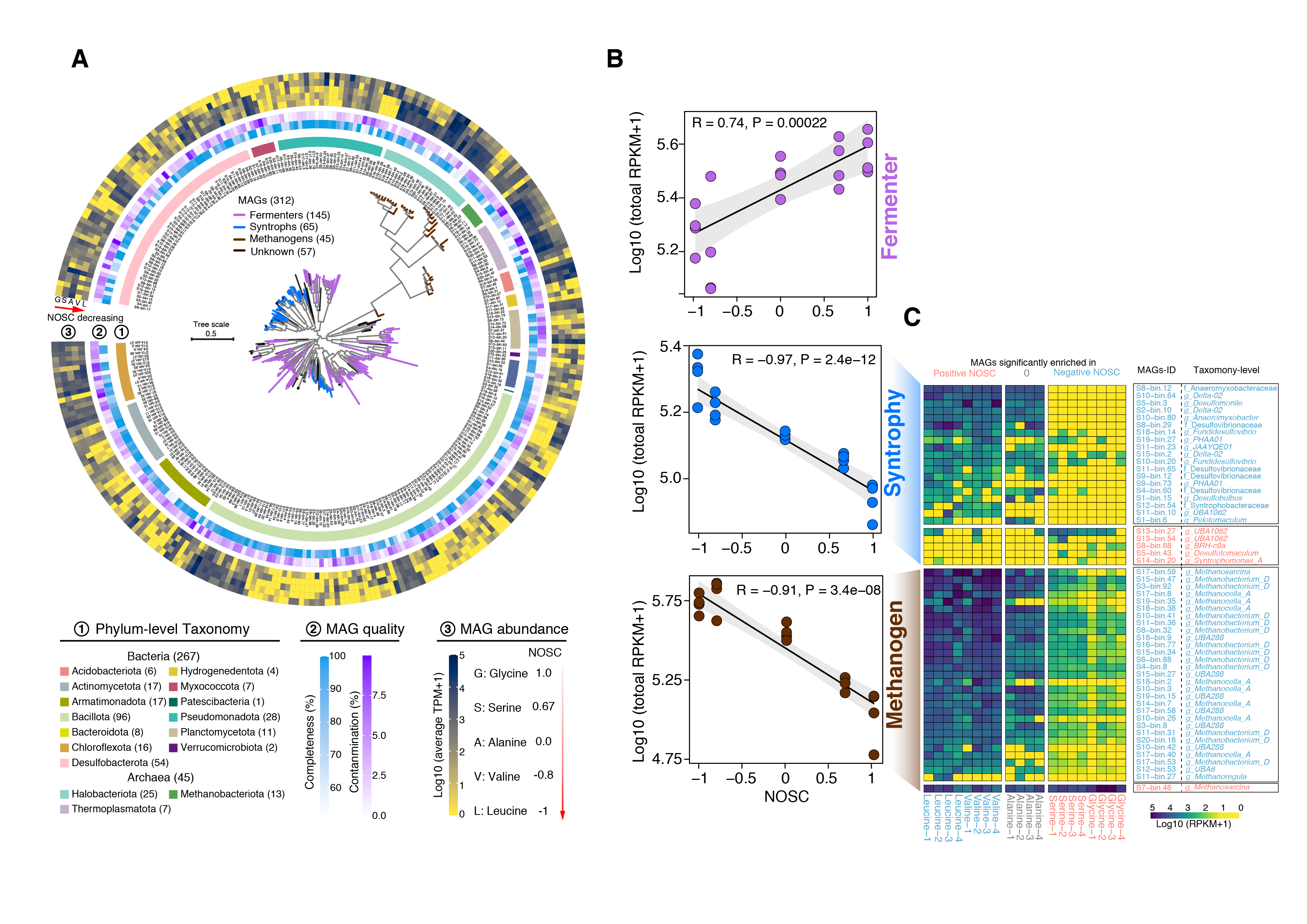

**Figure S5**: Summary of metagenome-assembled genomes (MAGs) with high or medium quality (completeness > 50% and contamination < 10%) obtained from enrichment cultures on five amino acids, including leucine, valine, alanine, serine and glycine. **A.** Phylogenetic analysis and relative abundances of the 312 MAGs obtained from enrichment cultures on five amino acids. **B.** Spearman correlations between relative abundances of fermentative, syntrophic or methanogenic MAGs and NOSC of five amino acids. **C.** Relative abundance of syntrophic and methanogenic microbial metagenome-assembled genomes (MAGs) differentially enriched in replicate microcosms amended with either negative NOSC amino acids (L-leucine and L-valine) or positive NOSC amino acids (L-serine and glycine). Each group of positive and negative NOSC enriched MAGs is sorted based on average Log2 fold-change (Log2FC). Mags-ID and Taxonomy-ID are given for each bin (See Supplemental Table S3).

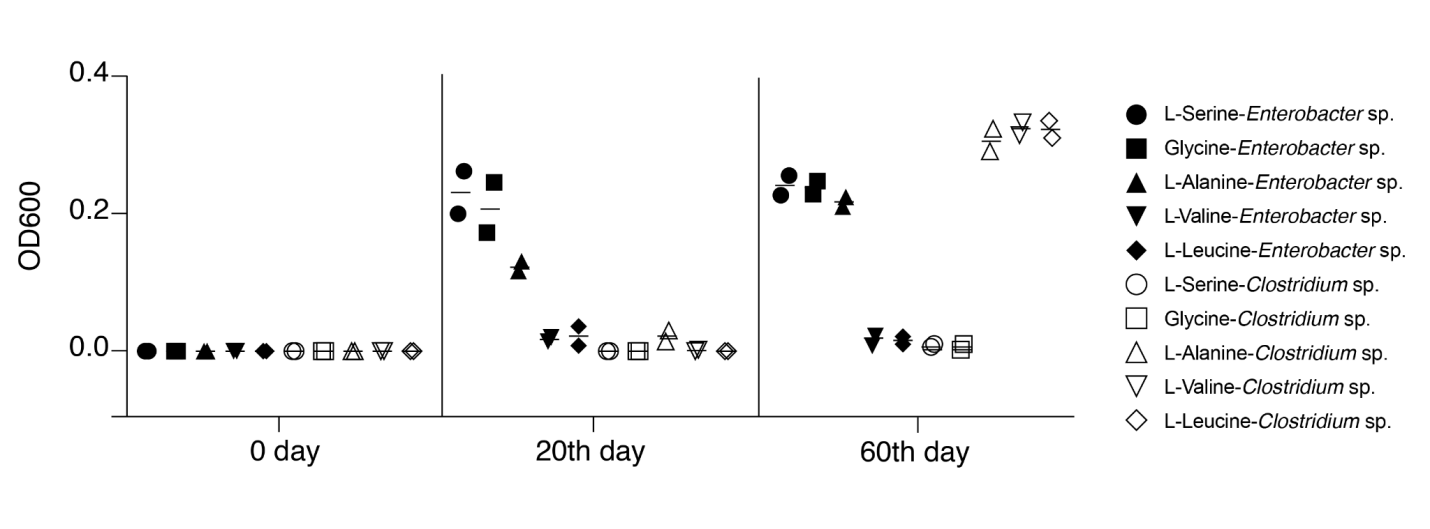

**Figure S6**: Comparison in growth yields of *Enterobacter* sp. and *Clostridium* sp. under anaerobic conditions across five amino acids.

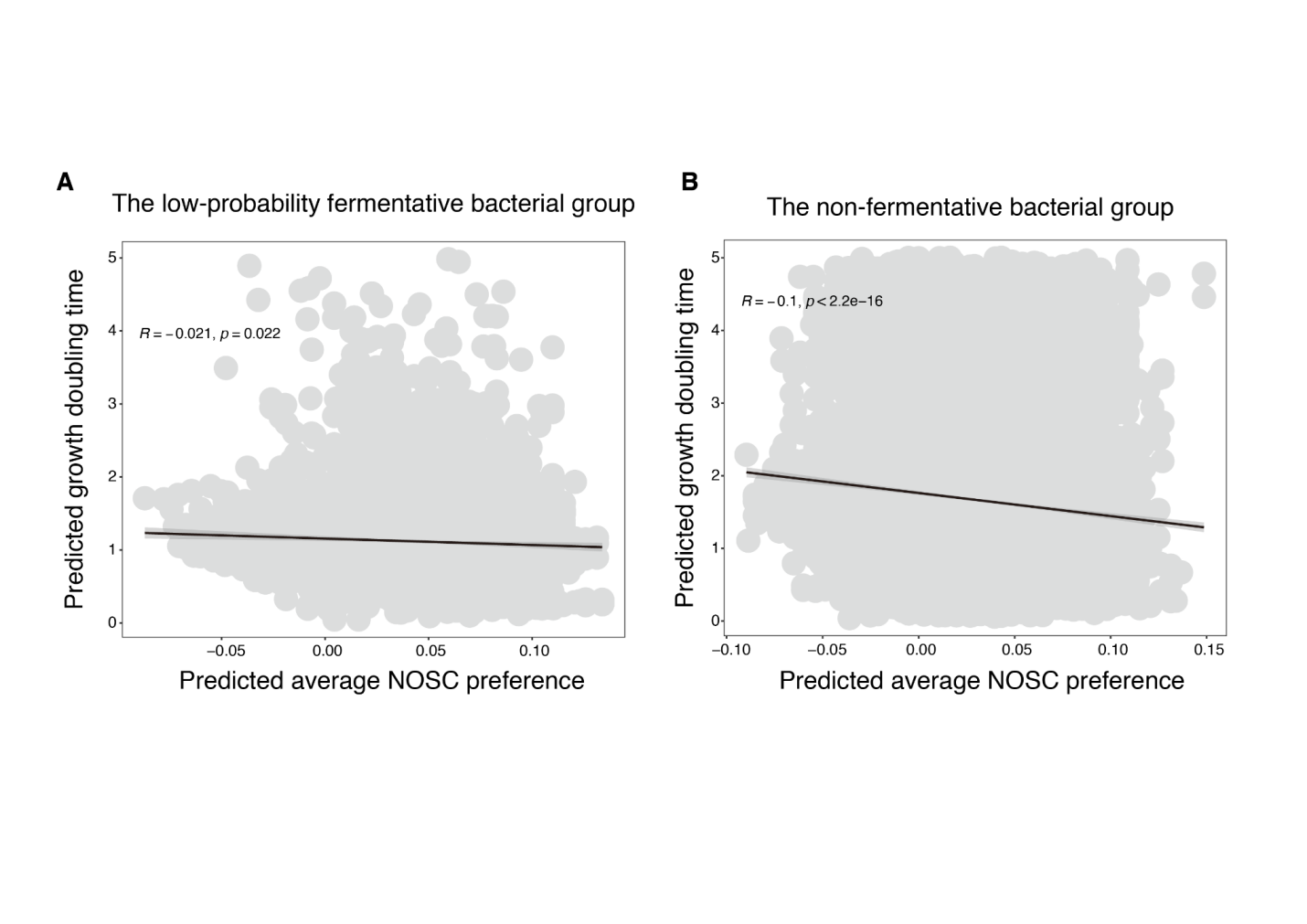

**Figure S7**: Spearman correlations between the predicted growth doubling time of 17016 bacterial genomes and their predicted average NOSC preference across 62 organic carbon sources from **A.** the low-probability fermentative bacterial group and **B.** the non-fermentative bacterial group.

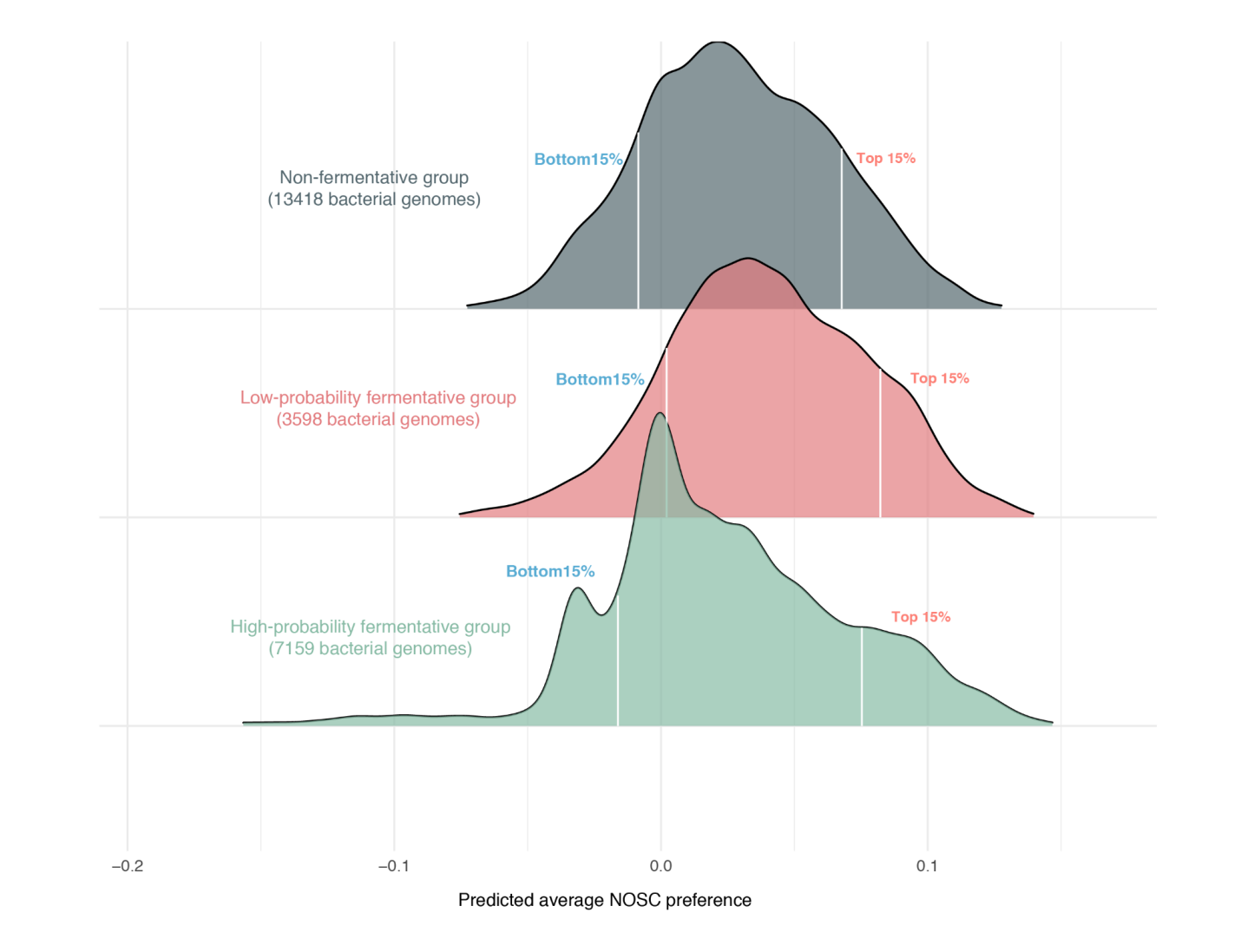

**Figure S8**: Distributions of the predicted average NOSC preference across 24,715 GTDB bacterial genomes from the high-probability fermentative group, the low-probability fermentative group and the non-fermentative bacterial group for 62 organic carbon compounds.
